# Desert Hedgehog-driven endothelium integrity is enhanced by Gas1 but negatively regulated by Cdon

**DOI:** 10.1101/2020.04.20.050542

**Authors:** Candice Chapouly, Pierre-Louis Hollier, Sarah Guimbal, Lauriane Cornuault, Alain-Pierre Gadeau, Marie-Ange Renault

## Abstract

Evidences accumulated within the past decades, identified Hedgehog (Hh) signaling as a new regulator of micro-vessel integrity. More specifically, we recently identified Desert Hedgehog (Dhh) as a downstream effector of Klf2 in endothelial cells (ECs).

**Objective:** The purpose of this study is to investigate whether Hh co-receptors Gas1 and Cdon may be used as therapeutic targets to modulate Dhh signaling in ECs.

**Methods and results:** We demonstrated that both Gas1 and Cdon are expressed in adult ECs and relied on either siRNAs or EC specific conditional KO mice to investigate their role. We found that Gas1 deficiency mainly photocopies Dhh deficiency especially by inducing VCAM-1 and ICAM-1 overexpression while Cdon deficiency has opposite effects by promoting endothelial junction integrity. At a molecular level, Cdon prevents Dhh binding to Ptch1 and thus acts a decoy receptor for Dhh, while Gas1 promotes Dhh binding to Smo and as a result potentiates Dhh effects. Since Cdon is overexpressed in ECs treated by inflammatory cytokines including TNFα and Il1β, we then tested whether Cdon inhibition would promote endothelium integrity in acute inflammatory conditions and found that both fibrinogen and IgG extravasation were decreased in association with an increased Cdh5 expression in the brain cortex of EC specific Cdon KO mice administered locally with Il1β.

**Conclusion:** Altogether these results demonstrate that Gas1 is a positive regulator of Dhh in ECs while Cdon is a negative regulator. Interestingly Cdon blocking molecules may then be used to promote endothelium integrity at least in inflammatory conditions.

## Introduction

Endothelium integrity is essential to vascular homeostasis, since a failure of this system represents a critical factor in cardiovascular and cerebrovascular disease pathogenesis. Indeed, the endothelium is involved in many physiological processes such as vascular permeability, vascular tone, blood coagulation and regulation as well as homing of immune cells to specific sites of the body. Conversely, endothelial dysfunction is associated with excessive vasoconstriction especially because of impaired endothelial nitric oxide (NO) production. Also, endothelial dysfunction is characterized by abnormal vascular leakage due to altered endothelial intercellular junctions. Finally, dysfunctional endothelial cells (ECs) acquire pro-inflammatory and pro-thrombotic phenotypes by expressing increased levels of adhesion and pro-thrombotic molecules such as vascular cell adhesion molecule-1 (VCAM-1) and intercellular adhesion molecule-1 (ICAM-1).

Evidences accumulated within the past decades, identified Hedgehog (Hh) signaling as a new regulator of micro-vessel integrity (Candice Chapouly et al. 2019a). For instance, Hh signaling was shown to promote blood brain barrier integrity and immune quiescence both in the setting of multiple sclerosis (Alvarez et al. 2011) and in the setting of stroke (Xia et al. 2013). Additionally we have shown that disruption of Hh signalling specifically in ECs induces blood-nerve barrier breakdown and peripheral nerve inflammation (C. Chapouly et al. 2016). While investigating molecular mechanisms underlying Hh dependent micro-vessel integrity, we found that endothelium “good function” depends on Desert Hedgehog (Dhh) expression by ECs themselves. More specifically, Dhh KO in ECs leads to the disruption of Cadherin-5 (Cdh5)/β-catenin interaction and spontaneous vascular leakage, an increased expression of adhesion molecules including VCAM-1 and ICAM-1 (Caradu et al. 2018) and increased angiogenic capabilities (Hollier et al. 2020). Importantly Dhh which is upregulated by blood flow and downregulated by inflammatory cytokines, appears to be a downstream effector of the master regulator of endothelial integrity Kruppel like factor 2 (Klf2) (Caradu et al. 2018).

The Hh family of morphogens which includes Sonic hedgehog (Shh), Indian hedgehog (Ihh) and Dhh, was identified nearly 4 decades ago in drosophila as crucial regulators of cell fate determination during embryogenesis (Nusslein-Volhard et Wieschaus 1980). The interaction of Hh proteins with their specific receptor Patched-1 (Ptch1) de-represses the transmembrane protein Smoothened (Smo), which activates downstream pathways, including the Hh canonical pathway leading to the activation of Gli family zinc finger (Gli) transcription factors and so-called Hh non canonical pathways, which are independent of Smo and/or Gli (Robbins, Fei, et Riobo 2012).

The Hh ligand binding to Ptch1 is regulated by several coreceptors. Among these, Cell adhesion molecule-related/downregulated by oncogenes (Cdon), Brother of Cdon (Boc) and Growth arrest specific 1 (Gas1) are suggested to promote Hh ligand interaction with Ptch1 while Hedgehog interacting protein (Hhip) inhibits it (Ramsbottom et Pownall 2016).

Cdon and Boc proteins are cell surface glycoproteins belonging to a subgroup of the Immunoglobulin superfamily of cell adhesion molecules, which also includes the Robo axon-guidance receptors. Their ectodomain respectively contains five and four Ig-like domains, followed by three type III fibronectin (FNIII) repeats (FNIII 1 to 3), a single trans-membrane domain and a divergent intracellular region of variable length (Sanchez-Arrones et al. 2012). Cdon was shown to interact with all of the three N-terminal active Hh peptides (Hh-N) through its third FNIII domain.

Gas1 was identified as one of six genes that were transcriptionally up‑regulated in NIH3T3 cells arrested in cell cycle upon serum starvation. Gas1 encodes a 45-kDa GPI-anchored cell surface protein that binds Shh-N with high affinity (Lee, Buttitta, et Fan 2001a).

The goal of the present study is to investigate whether Dhh-induced endothelial integrity depends on Hh co-receptors. This is essential to determine whether such co-receptors could be used as therapeutic target to enhance Dhh-induced signaling in ECs under pathological conditions.

## Methods

### Mice

Cdon Floxed (Cdon^Flox^) mice (Supplemental Figure I) were generated at the “Institut Clinique de la Souris” through the International Mouse Phenotyping Consortium (IMPC) from a vector generated by the European conditional mice mutagenesis program, EUCOMM. Gas1^tm3.1Fan^ (Gas1^Flox^) mice (Jin et al. 2015, 1) were kindly given by C.M. Fan and Tg(Cdh5-cre/ERT2)1Rha (Cdh5-CreERT2) mice (Azzoni et al. 2014) were a gift from RH. Adams.

Cdh5-Cre^ERT2^ mice were genotyped using the following primers: 5’-TAAAGATATCTCACGTACTGACGGTG-3’ and 5’-TCTCTGACCAGAGTCATCCTTAGC-3’ that amplify 493 bp of the Cre recombinase sequence. Cdon Floxed mice were genotyped using the following primers 5’-CTTCCCAGAGGGTGTGAGAGCAATG-3’ and 5’-GAACCAGTAGCATGCATGATGCTGG-3’ which amplifies a 385 bp fragment of the WT allele or a 494 bp fragment if the allele is floxed. Gas1 Floxed mice were genotyped using the following primers 5’-GAATCGAAGCGCCTGGACC-3’ and 5’-GGAAAACCGCACAGAAGAGGG-3’ which amplifies a 285 bp fragment of the WT allele or a 360 bp fragment if the allele is floxed.

Animal experiments were performed in accordance with the guidelines from Directive 2010/63/EU of the European Parliament on the protection of animals used for scientific purposes and approved by the local Animal Care and Use Committee of Bordeaux University.

The Cre recombinase in Cdh5-Cre^ERT2^ mice was activated by intraperitoneal injection of 1 mg tamoxifen for 5 consecutive days at 8 weeks of age. Mice were phenotyped 2 weeks later. Successful and specific activation of the Cre recombinase has been verified before (Caradu et al. 2018). Both males and females were used in equal proportions. At the end of experiments animal were sacrificed via cervical dislocation.

### Mouse corneal angiogenesis assay

Pellets were prepared as previously described (Kenyon et al. 1996). Briefly, 5 μg of VEGFA (Shenandoah biotechnology diluted in 10 μL sterile phosphate-buffered saline (PBS) was mixed with 2.5 mg sucrose octasulfate-aluminum complex (Sigma-Aldrich Co., St. Louis, MO, USA), and 10 μL of 12% hydron in ethanol was added. The suspension was deposited on a 400-μm nylon mesh (Sefar America Inc., Depew, NY, USA), then both sides of the mesh were covered with a thin layer of hydron and allowed to dry.

Female mice were anesthetized with an intraperitoneal (IP) injection of ketamine 100 mg/kg and xylazine 10 mg/kg. The eyes of the mice eyes were topically anesthetized with 0.5% Proparacaine™ or similar ophthalmic anesthetic. The globe of the eye was proptosed with jeweler’s forceps taking care to not damage the limbus vessel surrounding the base of the globe. Sterile saline was also be applied directly to each eye as needed during the procedure to prevent excessive drying of the cornea and to facilitate insertion of the pellet into the lamellar pocket of the eyes. Using an operating microscope, a central, intrasomal linear keratotomy was performed with a surgical blade parallel to the insertion of the lateral rectus muscle. Using a modified von greafe knife, a lamellar micro pocket was made toward the temporal limbus by ‘rocking’ the von greafe knife back and forth.

Hh containing or control pellet was placed on the cornea surface with jeweler’s forceps at the opening of the lamellar pocket. A drop of saline was applied directly to the pellet, and using the modified von greafe knife, the pellet was gently advanced to the temporal end of the pocket. Buprenorphine was given at a dose of 0.05 mg/kg subcutaneously on the day of surgery.

Nine days after pellet implantation, mice were sacrificed, and then eyes were harvested and fixed with 2% paraformaldehyde. Capillaries were stained with rat anti-mouse CD31 antibodies (BMA Biomedicals, Cat#T-2001), primary antibodies were visualized with Alexa 568–conjugated anti-rat antibodies (Invitrogen). Pictures were taken under 50x magnification. Angiogenesis was quantified as the CD31+surface area.

### In vivo permeability assessment (Miles assay)

The back of female mice was shaved. 72 hours, later mice were administered with 100 μL 1% Evans blue via retro orbital injection. Subsequently they were administered with 50 μL NaCl 0.9% containing or not 20 ng VEGFA (Shenandoah biotechnology) subcutaneously at 6 spots on their back. 30 minutes later mice were sacrificed, skin biopsy around each injection point were then harvested to quantify Evans blue extravasation. Evans blue dye was extracted from the skin by incubation at 65°C with formamide. The concentration of Evans blue dye extracted was determined spectrophotometrically at 620 nm with a reference at 740 nm. Buprenorphine was given at a dose of 0.05 mg/kg subcutaneously on the day of surgery.

### Ad-Il1β stereotaxic injections

Mice were anaesthetized using isoflurane and placed into a stereotactic frame (Stoelting). To prevent eye dryness, an ophthalmic *ointment* was applied at the ocular surface to maintain eye hydration during the time of surgery. The skull was shaved and the skin incised on 1 cm to expose the skull cap. Then, a hole was drilled into the cerebral cortex and 3 μL of an AdIL-1 (Horng et al. 2017) or AdDL70 control (AdCtrl), (10^7^ pfu) solution was microinjected at y=1 mm caudal to Bregma, x=2 mm, z=1.5 mm using a Hamilton syringe, into the cerebral cortex and infused for 3 minutes before removing the needle from the skull hole (Argaw et al. 2009). Mice received a subcutaneous injection of buprenorphine (0.05 mg/kg) 30 minutes before surgery and again 8 hours post-surgery to assure a constant analgesia during the procedure and postoperatively. Mice were sacrificed 7 days post-surgery. For histological assessment, brains were harvested and fixed in formalin for 3 hours before being incubated in 30% sucrose overnight and OCT embedded. Then, for each brain, the lesion area identified by the puncture site was cut into 7 μm thick sections.

### Immunostaining

Prior to staining, heart, brain, and lung tissues were fixed in methanol; paraffin embedded and cut into 7 μm thick sections. Whole mount corneas were fixed with 2.5% Formaline for 10 minutes and cultured cells were fixed with 10% formaline for 10 minutes.

Capillaries were identified using rat anti-mouse CD31 antibodies (BMA Biomedicals, Cat#T-2001). Neutrophils were stained with a rat anti-Ly6G (GR1) antibody (BD Pharmingen Inc, Cat#551459). Human Cdh5 was stained using mouse anti-human Cdh5 antibodies (Santa Cruz Biotechnology, Inc, Cat#sc-9989). Mouse Cdh5 was stained using goat anti-mouse Cdh5 antibodies (R&D systems, Cat# AF1002). Albumin and fibrinogen were stained using sheep anti-albumin antibodies (Abcam, Cat# ab8940) and rabbit anti-fibrinogen antibodies (Dako, Cat#A0080) respectively. Mouse IgGs were stained with Alexa Fluor 568 conjugated donkey anti-mouse IgG (Invitrogen, Cat#A-10037). Pan-leucocytes were identified using rat anti-mouse CD45 antibodies (BD Pharmingen Inc, Cat# 550539). CD11b+ microglia and macrophages were identified using rat anti-CD11b antibodies (ThermoFisher, cat#14-0112-82). GFAP was stained using rabbit anti-GFAP antibodies (ThermoFisher, Cat# OPA1-06100). Neurons were identified using anti-NeuN antibodies (Millipore, Cat# ABN78). Cdon was stained using goat anti-mouse Cdon antibodies (R&D systems, Cat#AF2429). Gas1 was stained using goat anti-human Gas1 antibodies (R&D systems, Cat# AF2636). Dhh was stained using mouse anti-Dhh antibodies (Santa Cruz Biotechnology, Inc, Cat#sc-271168). Ptch1 was stained using rabbit anti-Ptch1 antibodies (Abcam, Cat#ab53715). For immunofluorescence analyzes, primary antibodies were resolved with Alexa Fluor^®^–conjugated secondary polyclonal antibodies (Invitrogen, Cat# A-21206, A-21208, A-11077, A-11057, A-31573, A-10037) and nuclei were counterstained with DAPI (1/5000). For both immunohistochemical and immunofluorescence analyses, negative controls using secondary antibodies only were done to check for antibody specificity.

### Cell culture

*In vitro* experiments were performed using human umbilical vein endothelial cells (HUVECs) (Lonza), human dermal microvascular endothelial cells (HMVECs-D) (Lonza) or human brain microvascular Endothelial Cells (HBMECs) (Alphabioregen). HUVECs and HBMECs were cultured in endothelial basal medium-2 (EBM-2) supplemented with EGM™-2 BulletKits™ (Lonza). HMVECs-D were cultured in endothelial basal medium-2 (EBM-2) supplemented with EGM™-2 MV BulletKits™ (Lonza). Cell from passage 3 to passage 6 were used. Before any treatment cells were serum starved in 0.5% fetal bovine serum medium for 24 hours. HeLa ATCC^®^CCL-2™ cells were cultured in Roswell Park Memorial Institute medium (RPMI) supplemented with 10% fetal bovine serum.

### siRNA/Transfection

HUVECs were transfected with human Gas1 siRNA: rArGrCrArCrArUrUrUrCrUrUrArGrGrArUrUrArArGrGrGdTdC, human Cdon siRNA: rGrCrUrArUrArGrUrGrArCrArGrCrArArGrArGrCrUrCrCdTdC human Dhh siRNA: rArCrUrCrCrUrUrArArArGgArGrGrArCrUrArUrUrUrArGdCdC, human Ptch1 siRNA: rArGrArArArUrArCrCrCrArCrArGrCrArUrArGrUrGrArCdCdT: or universal scrambled negative control siRNA duplex (Origen) using JetPRIME™ transfection reagent (Polyplus Transfection), according to the manufacturer’s instructions.

### Plasmids/Transfection

The human Gas1 encoding vector, pcDNA3-Gas1 was kindly given by C.M. Fan (Lee, Buttitta, et Fan 2001a), the GFP tagged-mouse Cdon encoding vector, pCA-mCdonEGFP, was a gift from A. Okada (Okada et al. 2006), the myc-tagged human Ptch1, Ptch1-1B-myc was kindly given by R. Toftgard (Kogerman et al. 2002) and the human full length Dhh was previously described (Caradu et al. 2018). A myc tag was added by PCR at the N-terminal of human full length Dhh to generate the myc-tagged Dhh encoding vector.

HeLa cells were transfected using JetPRIME™ transfection reagent (Polyplus Transfection), according to the manufacturer’s instructions.

### Quantitative Reverse-Transcription Polymerase Chain Reaction (RT-PCR)

Following manufacturer’s instructions, RNAs were isolated and homogenized, from 3 × 10^5^ cells or from tissues previously snap-frozen in liquid nitrogen, using Tri Reagent^®^ (Molecular Research Center Inc). For quantitative RT-PCR analyzes, total RNA was reverse transcribed with M-MLV reverse transcriptase (Promega) and amplification was performed on a DNA Engine Opticon^®^2 (MJ Research Inc) using B-R SYBER^®^ Green SuperMix (Quanta Biosciences). Primer sequences are reported in Supplemental table I. The relative expression of each mRNA was calculated by the comparative threshold cycle method and normalized to β-actin mRNA expression.

### Immunoprecipitation/Western blot analysis

Prior to western blot analysis, Dhh, Ptch1 ou Smo were immunoprecipitated with mouse anti-Dhh antibodies (Santa Cruz Biotechnology, Cat# sc-271168), anti myc-tag antibodies (Millipore, Cat# 05-724) or mouse anti-Smo antibodies (Santa Cruz Biotechnology, Cat# sc-166685).

Expression of Cdon, Gas1, Dhh, Ptch1 and Smo were evaluated by SDS PAGE using goat anti-mouse Cdon antibodies (R&D systems, Cat#AF2429), goat anti-human Gas1 antibodies (R&D systems, Cat# AF2636), mouse anti-Dhh antibodies (Santa Cruz Biotechnology, Inc, Cat#sc-271168), rabbit anti-Ptch1 antibodies (Abcam, Cat#ab53715) and mouse anti-Smo antibodies (Santa Cruz Biotechnology, Cat# sc-166685) respectively.

Expression of human ICAM-1 and VCAM-1 expression were evaluated by SDS PAGE using mouse anti-human I-CAM1 antibodies (Santa Cruz Biotechnology, Cat#sc-8439) and rabbit anti-VCAM-1 (Abcam, Cat# ab134047) respectively.

Protein loading quantity was controlled using a monoclonal anti-α-tubulin antibody (Sigma). Secondary antibodies were from Invitrogen, Cat#A-21039, A-21084, A-21036). The signal was then revealed by using an Odyssey Infrared imager (LI-COR).

### In vitro permeability assay

100 000 cells were seeded in Transwell^®^ inserts. The day after, 0.5 mg/mL 70 kDa FITC-Dextran (Sigma) was added to the upper chamber. FITC fluorescence in the lower chamber was measured 20 minutes later.

### Migration assay

Cell migration was evaluated with a chemotaxis chamber (Neuro Probe, Inc., Gaithersburg, MD, USA). Briefly, a polycarbonate filter (8-μm pore size) (GE Infrastructure, Fairfield, CN, USA) was coated with a solution containing 0.2% gelatin (Sigma-Aldrich Co.) and inserted between the chambers, then 5×10^4^ cells per well were seeded in the upper chamber, and the lower chamber was filled with EBM-2 medium containing 0.5% FBS. Cells were incubated for 8 hours at 37°C then viewed under 20× magnification, and the number of cells that had migrated to the lower chamber were counted in 3 HPFs per well; migration was reported as the mean number of migrated cells per HPF. Each condition was assayed in triplicate and each experiment was performed at least three times.

### Methyl thiazolyl tetrazolium (MTT) cell proliferation Assay

5×10^3^ cells per well were seeded in a 96-well plate. At the indicated time points, 10 μL of 5 mg/mL MTT were added to each wells. Cells were incubated 3-4 hours at 37°C then culture medium was replaced by 100 μL DMSO. OD was read at 590 nm with a reference at 620 nm. Each condition included eight wells in each experiment and each experiment was performed at least three times.

### Statistics

Results are reported as mean ± SEM. Comparisons between groups were analyzed for significance with the non-parametric Mann-Whitney test or a one way ANOVA test followed by Bonferroni’s multiple comparison test (for than two groups) using GraphPad Prism v7.0 (GraphPad Inc, San Diego, Calif). Differences between groups were considered significant when p≤0.05 (*: p≤0.05; **: p≤0.01; ***: p≤0.001).

## Results

### ECs express Cdon, Gas1 and Hhip but not Boc

First, we searched for Cdon, Boc, Gas1 and Hhip expression in human EC from different origin, including HUVECs, HMVECs-D and HBMECs via RT-PCR. As shown in Figure 1A, human ECs express Hhip, Cdon, and Gas1 while they barely express Boc. Notably, Gas1 is not detected in HBMECs. Since the role of endothelial Hhip has already been reported in several papers (Agrawal, Kim, et Kwon 2017; Sekiguchi et al. 2012; Nie et al. 2016), we focused our investigations on Gas1 and Cdon and confirmed their endothelial expression via immunostaining (Figure 1B). Interestingly, while TNFα inhibits Gas1 mRNA expression in HUVECs (Supplemental Figure 2A), it increases Cdon mRNA expression (Supplemental Figure 2B). Moreover, TNFα-induced Cdon mRNA expression depends on NF-κB activity (Supplemental Figure 2C).

**Figure 1:**
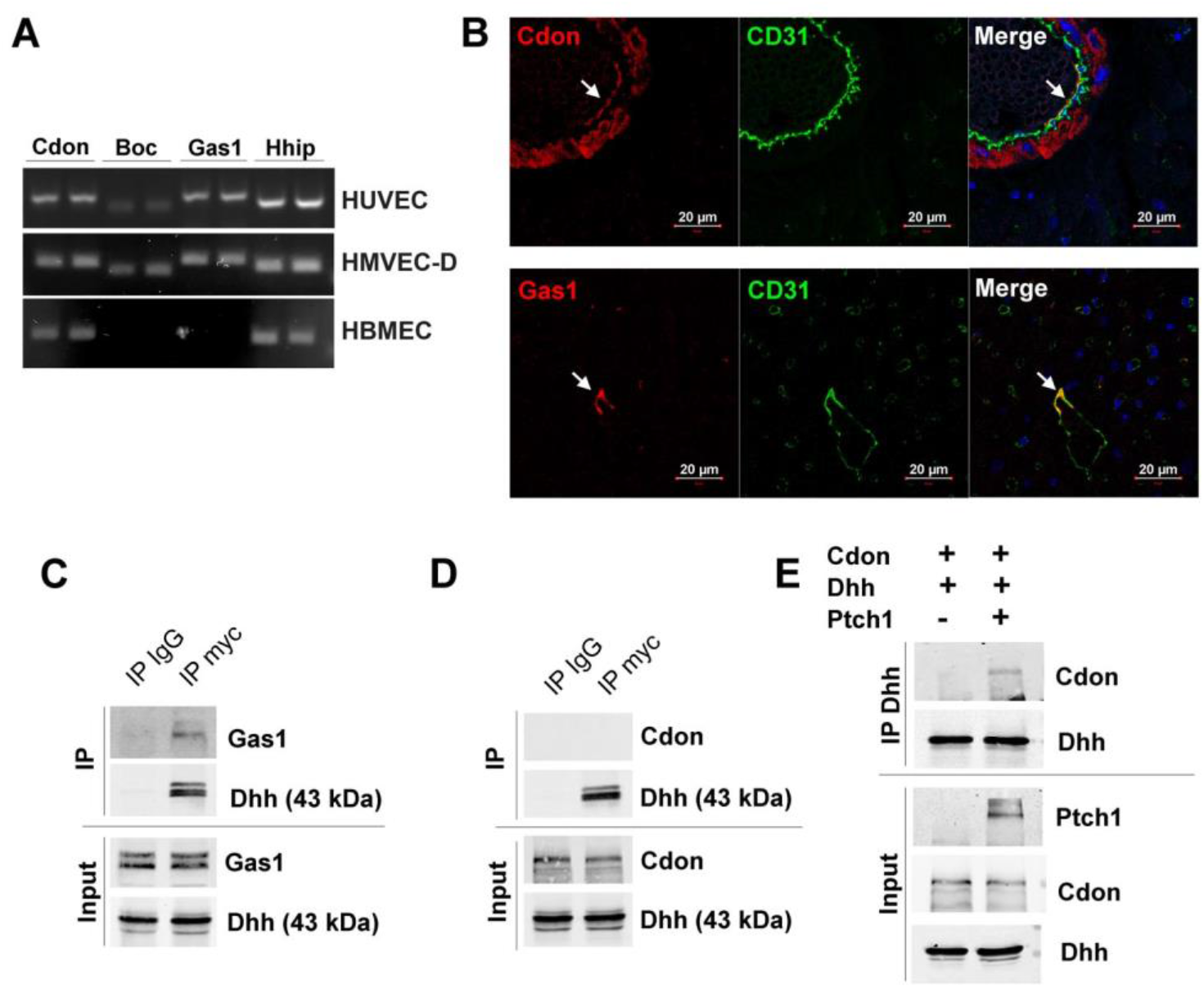
ECs express Cdon, Gas1 and Hhip. (**A**) Cdon, Boc, Gas1 and Hhip mRNA expression was evaluated via RT-PCR HUVECs, HMVECs-D and HBMECs. (**B**) Heart cross section from wild type mice were co-immunostained with anti-Cdon (in red) or anti-Gas1 (in red) antibodies together with anti-CD31 (in green) antibodies to identify Cdon or Gas1 expression in ECs respectively. (**C**) HeLa were co-transfected with Gas1 and myc-tagged Dhh encoding vectors. Gas1 interaction with Dhh was evaluated by co-immunoprecipitation assay. (**D**) HeLa were co-transfected with Cdon and myc-tagged Dhh encoding vectors. Cdon interaction with Dhh was evaluated by co-immunoprecipitation assay. (**E**) HeLa were co-transfected with Cdon, Ptch1 and Dhh encoding vectors. Cdon interaction with Dhh was evaluated by co-immunoprecipitation assay.

Next, we performed co-immunoprecipitation assays to verify that Dhh is able to bind these receptors. While we found that Dhh binds Gas1 directly (Figure 1C), Cdon alone cannot bind Dhh (Figure 1D). However, Dhh can bind Cdon in the presence of Ptch1 (Figure 1E). Consistently, Cdon co-localizes with Dhh only in the presence of Ptch1 while Gas1 co-localizes with Dhh both in the presence and absence of Ptch1 (Supplemental Figure 3)

With the aim to further investigate the role of Gas1 and Cdon in ECs we performed a series of *in vitro* and *in vivo* assays using siRNAs and EC-specific conditional KO mice respectively.

### Cdon promotes EC proliferation, migration and angiogenesis

The role of Gas1 and Cdon in angiogenesis was investigated using the mouse corneal angiogenesis assay. Mice deficient for Gas1 or Cdon expression in EC together with their respective control littermates were implanted with VEGFA containing pellets. While VEGFA-induced angiogenesis was not different in Gas1^ECKO^ mice from their control littermates (Figure 2A-B), VEGFA-induced angiogenesis was significantly-inhibited in Cdon^ECKO^ mice compared to their control littermates (Figure 2C-D). Consistently, *in vitro* experiments performed in HUVECs showed that both EC proliferation (Figure 2E) and VEGFA-induced EC migration (Figure 2F) were decreased after Cdon knock down (KD). Gas1 KD did not modify EC proliferation or VEGFA-induced migration. However, Gas1 KD did promote EC migration in the absence of VEGF (Figure 2E and 2F). This set of data demonstrates that Cdon is pro-angiogenic. Notably this effect works in the opposite direction to Dhh anti-angiogenic effect (Hollier et al. 2020).

**Figure 2:**
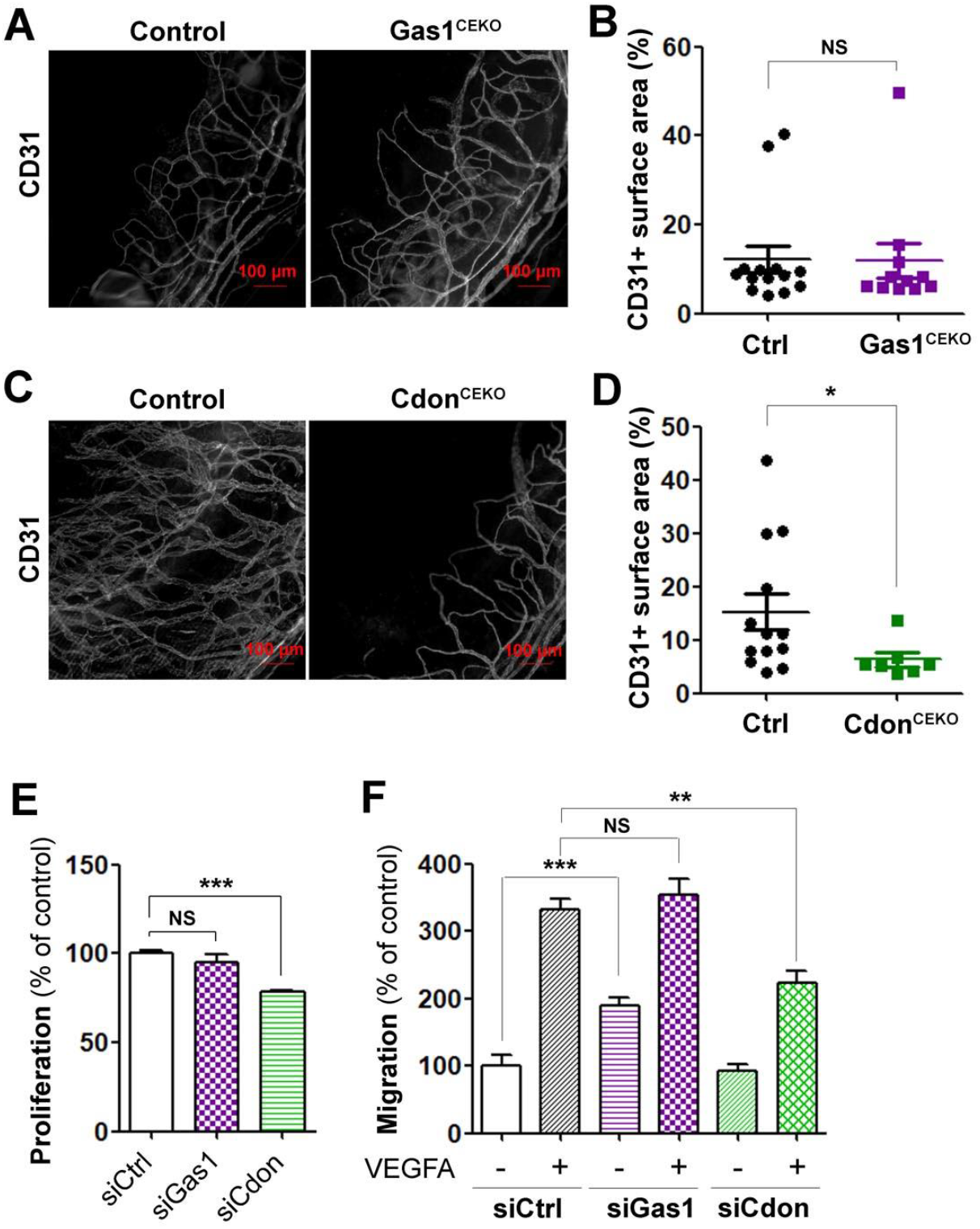
Cdon promotes EC migration, proliferation and angiogenesis. (**A-B)** VEGFA containing pellets were implanted in the corneas of Cdh5-Cre^ERT2^ Gas1^Flox/Flox^ (Gas1^ECKO^) and Gas1^Flox/Flox^ (control) mice 2 week after they were administered with tamoxifen (n=14 and 11 corneas respectively). (**A**) Whole mount corneas were immunostained with anti-CD31 antibodies to identify blood vessels. Representative pictures are shown. (B) Angiogenesis was quantified as the percentage of CD31+ surface area. (**C-D)** VEGFA containing pellets were implanted in the corneas of Cdh5-Cre^ERT2^ Cdon^Flox/Flox^ (Cdon^ECKO^) and Cdon^Flox/Flox^ (control) mice 2 week after they were administered with tamoxifen (n=13 and 7 corneas respectively). (**C**) Whole mount corneas were immunostained with anti-CD31 antibodies to identify blood vessels. Representative pictures are shown. (**D**) Angiogenesis was quantified as the percentage of CD31+ surface area. (**E-F**) HUVECs were transfected with Gas1, Cdon or control siRNAs. (**E**) Cells proliferation was assessed using MTT. The experiment was repeated 3 times, each experiment included n=8 wells/conditions. (**F**) Cell migration was assessed in a chemotaxis chamber in the presence or not of 50 ng/mL VEGFA. The experiment was repeated 3 times, each experiment included n=4 wells/conditions. *: p≤0.05; **: p≤0.01; ***: p≤0.001. NS: not significant. Mann Whitney test or one way ANOVA followed by Bonferroni’s multiple comparisons test.

### Gas1 prevent EC activation and LPS-induced neutrophil recruitment

To investigate the role of Gas1 and Cdon in regulating EC immune quiescence, HUVECs were transfected with Gas1, Cdon or control siRNAs. VCAM-1 and ICAM-1 expression was measured via both RT-qPCR and western blot analyses. While Gas1 KD significantly increased both VCAM-1 and ICAM-1 expression (Figure 3A-C), Cdon KD did not (Figure 3A-C). These results were confirmed in TNFα treated cells (Figure 3D-E). Finally, to assess the functional consequences of EC activation *in vivo*, we quantified neutrophils recruitment in the lungs of mice that were administered with LPS. As expected, neutrophils density in the lung of Gas1^ECKO^ mice was significantly increased (Figure 3F-G) compared to their control littermates while we found no difference between Cdon^ECKO^ mice and control littermates (Figure 3H-I).

**Figure 3:**
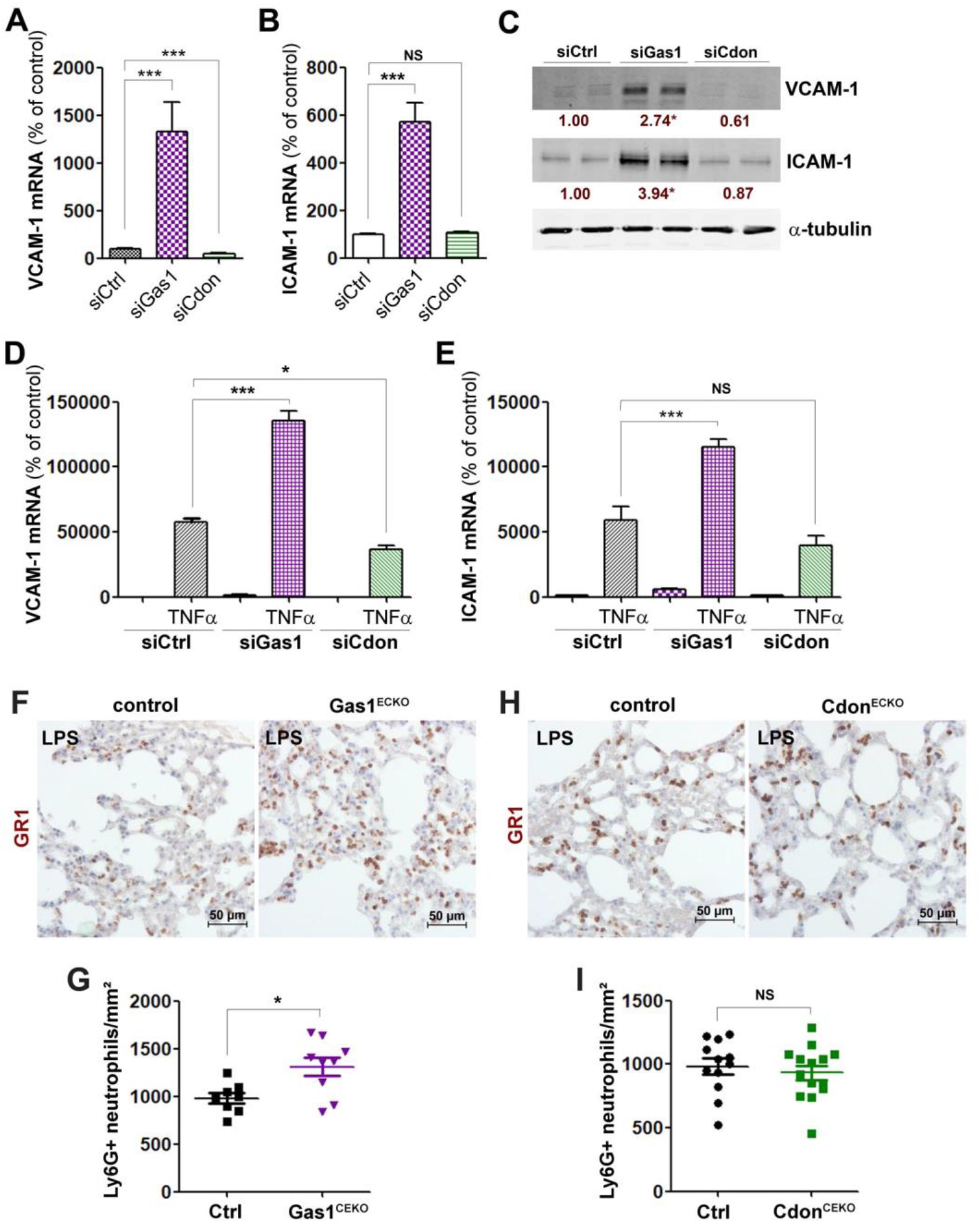
Gas1 prevents EC activation and neutrophils recruitment. (**A-C**) HUVECs were transfected with Gas1, Cdon or control siRNAs. VCAM-1 (**A**) and ICAM-1 (**B**) mRNA expression was quantified via RT-qPCR. The experiment was repeated 3 times, each experiment included triplicates. (**C**) VCAM-1 and ICAM-1 protein expression was quantified by western blot analysis the experiment was repeated at least 3 times, each experiment included duplicates. (**D-E**) HUVECs were transfected with Gas1, Cdon or control siRNAs and then treated or not with 10 ng/mL TNFα for 6 hours. VCAM-1 (**D**) and ICAM-1 (**E**) mRNA expression was quantified via RT-qPCR. The experiment was repeated 3 times, each experiment included triplicates. (**F-G**) Cdh5-Cre^ERT2^ Gas1^Flox/Flox^ (Gas1^ECKO^) and Gas1^Flox/Flox^ (control) mice were treated with 10 mg/kg LPS and sacrificed 6 hours later. Lung sections were immunostained with anti-ICAM-1 (**F**) Lung sections were immunostained with anti-Ly6G (GR1) antibodies to identify neutrophils. (**G**) Neutrophil infiltration was quantified as the number of Ly6G+ cells/mm² (n=9 and 8 respectively). (**H-I**) Cdh5-Cre^ERT2^ Cdon^Flox/Flox^ (Cdon^ECKO^) and Cdon^Flox/Flox^ (control) mice were treated with 10 mg/kg LPS and sacrificed 6 hours later. (**H**) Lung sections were immunostained with anti-Ly6G (GR1) antibodies to identify neutrophils. (**I**) Neutrophil infiltration was quantified as the number of Ly6G+ cells/mm² (n=14 and 12 respectively). *: p≤0.05; ***: p≤0.001. NS: not significant. Mann Whitney test or one way ANOVA followed by Bonferroni’s multiple comparisons test.

Altogether these data demonstrate that Gas1 prevents EC activation similarly to Dhh (Caradu et al. 2018). On the contrary, Cdon does not seem to participate in the regulation of EC activation.

### Cdon disrupts adherens junction integrity

The role of Gas1 and Cdon in controlling endothelial intercellular junction integrity was first investigated *in vitro*. HUVECs were transfected with Gas1, Cdon or control siRNAs. Adherens junction integrity was quantified after Cdh5 immunostaining (Figure 4A) and endothelium permeability using Transwells. We found that Gas1 KD disrupts Cdh5-dependent junction integrity (Cdh5 junctions acquire a Zigzag phenotype) (Figure 4B) while Cdon KD prevents EC permeability (Figure 4C). Gas1 KD did not show any effects in the permeability assay test suggesting a very mild effect of Gas1 on adherens junction integrity. Consistently, in the Miles assay *in vivo,* VEGFA-induced vascular permeability was not different between Gas1^ECKO^ and control mice (Figure 4D) while it was significantly decreased in the absence of endothelial Cdon (Figure 4E).

**Figure 4:**
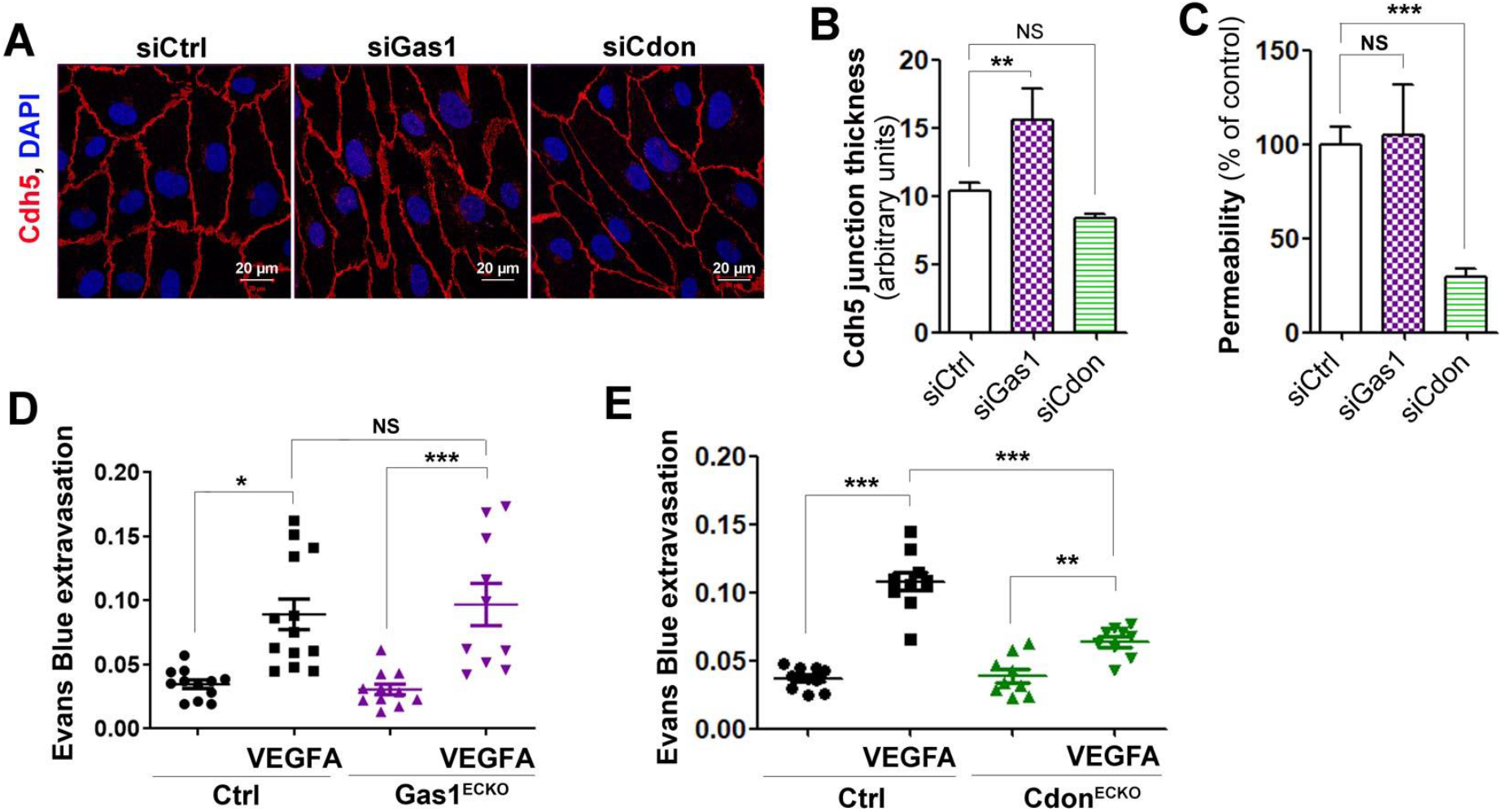
Gas1 promotes adherens junction integrity while Cdon disrupts it. (**A-C**) HUVECs were transfected with Gas1, Cdon or control siRNAs. (**A**) Cdh5 localization was evaluated by immunofluorescent staining (in red) of a confluent cell monolayer and (**B**) quantified as the mean junction thickness using Image J software. The experiment was repeated at least 4 times. (**C**) Endothelial monolayer permeability to 70 kDa FITC-Dextran was assessed using Transwells. The experiment was repeated 3 times, each experiment included triplicates. (**D**) VEGFA-induced permeability was assessed in both Cdh5-Cre^ERT2^ Gas1^Flox/Flox^ (Gas1^ECKO^) and Gas1^Flox/Flox^ (control) mice using the Miles assay (n=10 and 14 mice respectively). (**E**) VEGFA-induced permeability was assessed in both Cdh5-Cre^ERT2^; Cdon^Flox/Flox^ (Cdon^ECKO^) and Cdon^Flox/Flox^ (control) mice using the Miles assay (n=9 and 10 mice respectively). *: p≤0.05; **: p≤0.01; ***: p≤0.001. NS: not significant. One way ANOVA followed by Bonferroni’s multiple comparisons test.

This last set of data demonstrate that Cdon strongly increases vascular permeability unlike Dhh (Hollier et al. 2020; Caradu et al. 2018) and that Gas 1 may slightly modify Cdh5 junction organization without functional consequences.

To conclude on this first set of results, Gas1 may promote Hh signaling in ECs since Gas1 KD mostly recapitulates the effects of Dhh KD. On the contrary, Cdon most likely inhibits Hh signaling since Cdon KD induces opposite effect to those of Dhh KD. Notably, Cdon has been previously identified as a Hh decoy receptor in the zebrafish optic vesicle (Cardozo et al. 2014).

Therefore, in the second part of this study, we chose to perform a series of experiments aiming to investigate whether and how Gas1 and Cdon modulate Hh signaling in ECs.

### Gas1 promotes Dhh interaction with Smo while Cdon prevents Dhh interaction with Ptch1

First we tested whether Gas1 or Cdon modulate Dhh interaction with Ptch1 and Smo. Notably, Smo has been recently shown to be a receptor for Hh ligands especially in the case of cell autonomous signaling (Casillas et Roelink 2018). Interestingly, we found that Gas1 prevents Dhh interaction with Ptch1 but promotes Dhh interaction with Smo. On the contrary, Cdon prevents Dhh interaction with Ptch1 but does not modify Dhh interaction with Smo (Figure 5A-B).

**Figure 5:**
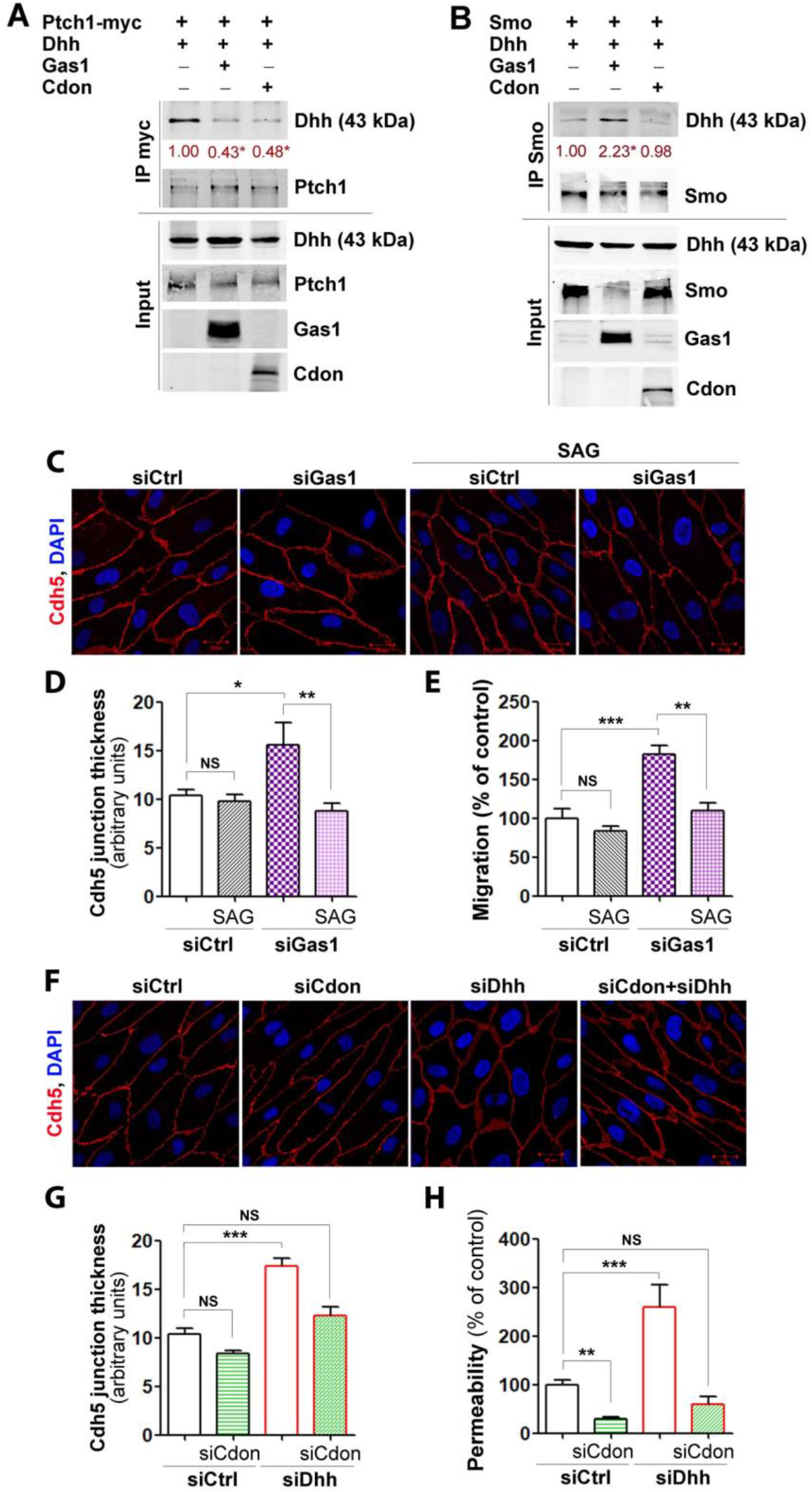
Gas1 promotes Dhh binding to Smo while Cdon prevents Dhh binding to Ptch1. (**A**) HeLa were co-transfected by Ptch1-myc and Dhh encoding plasmids together with or without Gas1 or Cdon encoding plasmids. Dhh interaction with Ptch1 was assessed by co-immunoprecipitation assay. The experiment was repeated 4 times. (**B**) HeLa were co-transfected Smo and Dhh encoding plasmids together with or without Gas1 or Cdon encoding plasmids. Dhh interaction with Smo was assessed by co-immunoprecipitation assay. The experiment was repeated 4 times. (**C-E**) HUVECs were transfected with Gas1 or control siRNAs, then treated or not with 100 nM SAG. (**C**) Cdh5 localization was evaluated by immunofluorescent staining (in red) of a confluent cell monolayer and (**D**) quantified as the mean junction thickness using Image J software. The experiment was repeated at least 4 times. (**E**) Cell migration was assessed in a chemotaxis chamber. The experiment was repeated 3 times, each experiment included n=4 wells/conditions. (**F-I**) HUVECs were co-transfected with Cdon or control siRNAs together with or without Dhh siRNAs. (**F**) Cdh5 localization was evaluated by immunofluorescent staining (in red) of a confluent cell monolayer and (**G**) quantified as the mean junction thickness using Image J software. The experiment was repeated at least 4 times. (**H**) Endothelial monolayer permeability to 70 kDa FITC-Dextran was assessed using Transwells. The experiment was repeated 3 times, each experiment included triplicates. The experiment was repeated 3 times, each experiment included n=4 wells/conditions. *: p≤0.05; **: p≤0.01; ***: p≤0.001. NS: not significant. One way ANOVA followed by Bonferroni’s multiple comparisons test.

Since Gas1 KD phenocopies most features of Dhh deficiency, we tested whether Gas1 effects on endothelial adherens junction’s integrity and migration depend on Hh signaling. To do so, we performed rescue experiments. HUVECs were either transfected with Gas1 or control siRNAs and then treated or not with the Smo agonist SAG. As shown in Figure 5C and 5D, Gas1 KD-induced Cdh5 junction thickening was prevented in the presence of SAG. Similarly, Gas1 KD failed to induce EC migration in the presence of SAG (Figure 5E). However, SAG had no effect on Gas1 KD-induced VCAM-1 and ICAM-1 (Supplemental Figure 4A and 4B). Notably, VCAM-1 and ICAM-1 seem to be downstream of Ptch1 rather than Smo since Ptch1 KD is sufficient to increase their expression (Supplemental Figure 4C and 4D).

Because Cdon has opposite effects to Dhh ones, we hypothesized that Cdon is a decoy receptor for Dhh at the surface of EC and thus tested whether siCdon-induced effects are prevented in the absence of Dhh. HUVECs were transfected with Cdon siRNAs alone or in combination with Dhh siRNAs. While siCdon alone decreased adherens junction thickness and endothelium permeability, in the siCdon + siDhh condition (Figure 5 F-H), effects were no longer significant confirming our hypothesis.

### Cdon deficiency at the endothelium prevents blood-brain barrier opening in the setting of acute inflammation

Finally, since Cdon appears to act as a negative regulator of Dhh-induced signaling in ECs, we hypothesized that blocking Cdon may promote Dhh-induced signaling in ECs and subsequently promote maintenance of endothelium integrity in pathological conditions.

To test such hypothesis, we administered adenoviruses encoding Il1β locally in the cortex of both Cdon^ECKO^ mice and their control littermate to induce acute brain inflammation and BBB breakdown. Notably, Cdon expression is significantly increased upon Il1β treatment in both HUVECs and HBMECs (Figure 6A-B). In accordance with our hypothesis, endothelial adherens junctions were preserved in the absence of Cdon, as attested by an increased Cdh5 expression in the cortical lesion area of Cdon^ECKO^ mice injected with Il1β, compared to control littermates (Figure 6D-E). Consistently, both fibrinogen and IgG extravasation were decreased (Figure 6D, F-G). BBB tightness in Cdon^ECKO^ mice was associated with a decreased leucocyte infiltration, a decreased microglia and astrocyte activation and finally with an increased neuron survival (Supplemental figure 5A-E).

**Figure 6:**
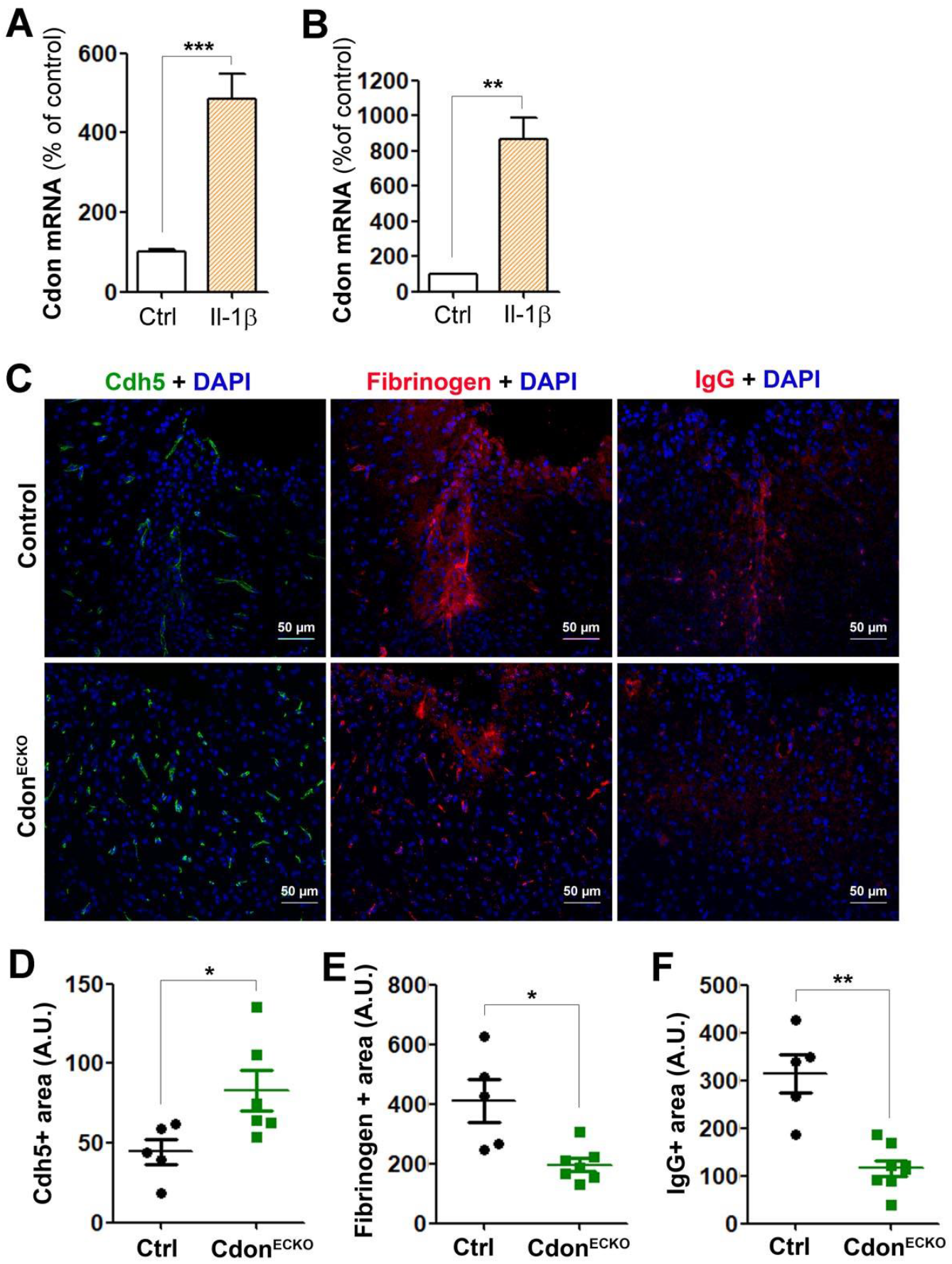
Cdon deficiency in ECs prevents Il1β-induced BBB disruption. HUVECs (**A**) and HBMECs (**B**) were treated or not with 10 ng/mL Il1β for 6 hours. Cdon mRNA expression was quantified by via RT-qPCR. Experiments were repeated 3 times, each experiment included triplicates. (**C-F**) Both Cdh5-Cre^ERT2^; Cdon^Flox/Flox^ (Cdon^ECKO^) and Cdon^Flox/Flox^ (control) mice were administered in the cerebral cortex with adenoviruses encoding Il1β (n=7 and 5 mice respectively). Mice were sacrificed 7 days later. (**C**) Brain sagittal sections were immunostained with anti-Cdh5 (in green), anti-Fibrinogen (in red) or anti-IgG (in red) antibodies. Representative confocal images are shown. (**D**) Cdh5 expression was quantified as the Cdh5+ surface area. (**E**) Fibrinogen extravasation was quantified as the fibrinogen+ surface area. (**F**) IgG extravasation was quantified as the fibrinogen+ surface area. *: p≤0.05; **: p≤0.01; ***: p≤0.001. Mann Withney test.

These last results demonstrate that blocking Cdon might indeed be a working therapeutic strategy to preserve endothelium integrity in pathological setting such as acute neuro-inflammation.

### Cdon blocking antibodies may be used as a therapeutic tool to maintain endothelial junctions in the setting of inflammation

We then tested whether Cdon antibodies may be used as a therapeutic tool to block Dhh binding to Cdon and improve endothelial integrity. To do so, HUVECs were treated or not with TNFα, in the presence or not of Cdon blocking antibodies. As shown in Figure 7A and 7B, TNFα-induced Cdh5 junction thickening is prevented in the presence of Cdon blocking antibodies.

**Figure 7:**
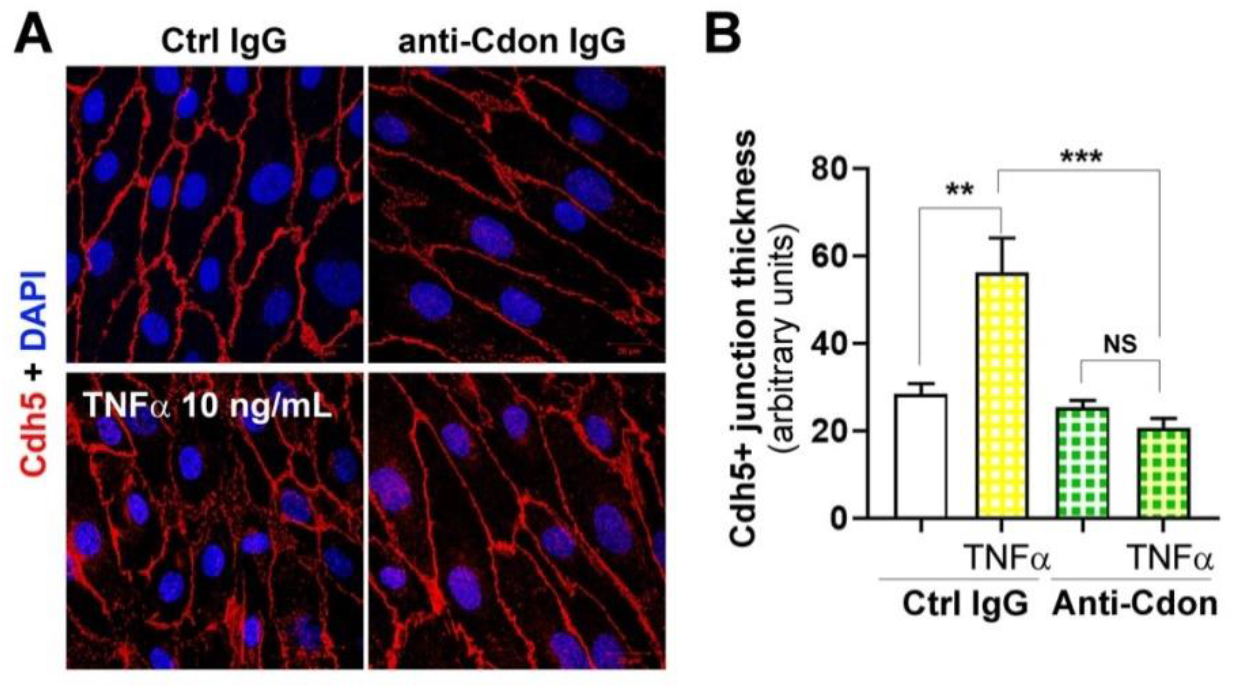
Cdon blocking antibodies can be used to promote endothelium integrity. HUVECs were treated or not with 10 ng/mL TNFα in the presence of 1.5 μg/mL Cdon blocking antibodies or 1.5 μg/mL unspecific IgGs. (**A**) Cdh5 localization was evaluated by immunofluorescent staining (in red) of a confluent cell monolayer and (**B**) quantified as the mean junction thickness using Image J software. The experiment was repeated at least 4 times. **: p≤0.01; ***: p≤0.001. NS: not significant. One way ANOVA followed by Bonferroni’s multiple comparisons test.

## Discussion

Hh signaling has been described to be regulated by several co-receptors including Hhip, Boc, Cdon and Gas1 especially in the setting of embryogenesis (Allen et al. 2011). The purpose of the present study was to investigate the role of Gas1 and Cdon in ECs in adults. Importantly, Hh signaling in ECs is original by several aspects. First, it exclusively involves non canonical signaling (Renault et al. 2010; Chinchilla et al. 2010), second, it is activated by full length unprocessed Dhh (FL-Dhh) (Hollier et al. 2020) and third, it occurs cell autonomously (Caradu et al. 2018). It is important to have in mind that full length unprocessed Hh ligands, may not only bind Ptch1 but also Smo directly (Casillas et Roelink 2018). In this particular setting, the present study demonstrates that Cdon prevents FL-Dhh binding to Ptch1. Gas1 also prevents FL-Dhh binding to Ptch1 but promotes FL-Dhh binding to Smo. By doing so, Cdon mainly acts as a negative regulator of FL-Dhh and destabilizes EC junctions to promote angiogenesis while Gas 1 is a positive regulator of FL-Dhh which prevents EC activation (Supplemental Figure 6).

Cdon, Gas1 and Boc are typically believed to be positive regulators of Hh signaling (Ramsbottom et Pownall 2016) in line with the fact that Gas1, Cdon and Boc were shown to be equally capable of promoting Shh signaling during neural patterning since overexpression of any individual component results in ectopic ventral cell fate specification (Allen et al. 2011). Additionally, while genetic removal of Gas1, Cdon or Boc individually has only modest effects on Shh signaling, removal of any two components results in significantly reduced Shh-dependent ventral neural patterning (Allen et al. 2011). However, conflicting results have been published: Gas1 was first shown to bind Shh in 2001. However, it was first suggested to reduce the availability of active Shh in the somite based on ectopic expression studies (Lee, Buttitta, et Fan 2001b). In 2007, experiments using Gas1 deficient mice revealed, on the contrary, that Gas1 is a positive regulator of Shh signaling and facilitates Shh low level effects (Martinelli et Fan 2007). Similarly, Cdon was shown to positively regulate Shh-induced signaling especially in the developing brain (Tenzen et al. 2006; Zhang et al. 2006) while it was more recently shown to act as a Hh decoy receptor during proximal-distal patterning of the optic vesicle (Cardozo et al. 2014). Whether Gas1 and Cdon are positive or negative regulators of Hh signaling may then most likely depend on the type of ligand and cell type involved.

Hh signaling in ECs is still far from being fully understood (Candice Chapouly et al. 2019b). We have previously shown that Dhh prevents EC activation by downregulating VCAM-1 and ICAM-1 and protects adherens junction integrity by promoting Cdh5 interaction with β-catenin (Caradu et al. 2018). The present study suggests that Dhh regulates EC activation and EC junction integrity via distinct pathways. Indeed, while Cdon mainly affects Dhh regulation of endothelial junctions, Gas1 mainly regulates Dhh regulation of EC immune quiescence. In both cases, a dialogue between Ptch1 and Smo seems to be involved, since Cdon modulates Dhh interaction with Ptch1 to regulate EC junctions, while we previously found that Cdh5 junction integrity depends on Smo (Hollier et al. 2020). Similarly, Gas1 promotes Dhh binding with Smo to prevent EC activation, while we found that Ptch1 KD is sufficient to increase VCAM-1 and ICAM-1 expression in ECs. We then hypothesized that both dialogues going from Ptch1 to Smo and Smo to Ptch1 exist (Supplemental Figure 6) based on the reciprocal regulation of Ptch1 and Smo by Smurf family of E3 ubiquitin ligases (Li et al. 2018).

Finally, the main goal of this study was to investigate whether Hh co-receptors may be used to modulate Hh signaling in ECs for therapeutical purposes. By identifying Cdon as a negative regulator of Dhh in ECs, and by demonstrating that Cdon KO prevents BBB opening in the setting of brain inflammation, the present study offer the possibility of using Cdon blocking molecules including blocking antibodies (Figure 7) as therapeutic tools to preserve endothelial integrity at least in the setting of inflammation. Notably, inflammatory cytokines including TNFα and Il1β increase Cdon expression in ECs.

## Acknowledgment

We thank Carnegie for proving with the Gas1 Floxed mice. We thank Annabel Reynaud, Sylvain Grolleau, and Maxime David for their technical help. We thank Christelle Boullé for administrative assistance.

This study was supported by grants from the Fondation de France (Appel d’Offre Recherche sur les maladies Cardiovasculaires 2013 and 2018), the Fondation pour la Recherche Médicale (équipe FRM) and the Fondation ARSEP pour la recherche sur la sclérose en plaques. Also this study was funded by a Marie Skłodowska-Curie Actions (MSCA-IF-2019) from the European concil. Finally, this study was co-funded by the “Institut National de la Santé et de la Recherche Médicale” and by the University of Bordeaux.

## Author contributions

C.C. conducted experiments, acquired data, analyzed data. P.-L. H., S.G. and L.C. conducted experiments, acquired data. A.-P. G. critically revised the manuscript. M.-A. R. designed research studies, conducted experiments, acquired data, analyzed data, providing reagents, and wrote the manuscript

## References

Agrawal, V., D. Y. Kim, et Y. G. Kwon. 2017. « Hhip Regulates Tumor-Stroma-Mediated Upregulation of Tumor Angiogenesis ». Exp Mol Med 49 (1): e289. https://doi.org/10.1038/emm.2016.139 emm2016139 [pii].

Alvarez, J. I., A. Dodelet-Devillers, H. Kebir, I. Ifergan, P. J. Fabre, S. Terouz, M. Sabbagh, et al. 2011. « The Hedgehog Pathway Promotes Blood-Brain Barrier Integrity and CNS Immune Quiescence ». Science 334 (6063): 1727–31.

Argaw, Azeb Tadesse, Blake T. Gurfein, Yueting Zhang, Andleeb Zameer, et Gareth R. John. 2009. « VEGF-Mediated Disruption of Endothelial CLN-5 Promotes Blood-Brain Barrier Breakdown ». Proceedings of the National Academy of Sciences of the United States of America 106 (6): 1977–82. https://doi.org/10.1073/pnas.0808698106.

Azzoni, E., V. Conti, L. Campana, A. Dellavalle, R. H. Adams, G. Cossu, et S. Brunelli. 2014. « Hemogenic Endothelium Generates Mesoangioblasts That Contribute to Several Mesodermal Lineages in Vivo ». Development 141 (9): 1821–34. https://doi.org/10.1242/dev.103242141/9/1821 [pii].

Caradu, Caroline, Thierry Couffinhal, Candice Chapouly, Sarah Guimbal, Pierre-Louis Hollier, Eric Ducasse, Alessandra Bura-Rivière, Mathilde Dubois, Alain-Pierre Gadeau, et Marie-Ange Renault. 2018. « Restoring Endothelial Function by Targeting Desert Hedgehog Downstream of Klf2 Improves Critical Limb Ischemia in Adults ». Circulation Research 123 (9): 1053–65. https://doi.org/10.1161/CIRCRESAHA.118.313177.

Cardozo, M. J., L. Sanchez-Arrones, A. Sandonis, C. Sanchez-Camacho, G. Gestri, S. W. Wilson, I. Guerrero, et P. Bovolenta. 2014. « Cdon Acts as a Hedgehog Decoy Receptor during Proximal-Distal Patterning of the Optic Vesicle ». Nat Commun 5: 4272.

Casillas, Catalina, et Henk Roelink. 2018. « Gain-of-Function Shh Mutants Activate Smo Cell-Autonomously Independent of Ptch1/2 Function ». Mechanisms of Development 153: 30–41. https://doi.org/10.1016/j.mod.2018.08.009.

Chapouly, C., Q. Yao, S. Vandierdonck, F. Larrieu-Lahargue, J. N. Mariani, A. P. Gadeau, et M. A. Renault. 2016. « Impaired Hedgehog Signalling-Induced Endothelial Dysfunction Is Sufficient to Induce Neuropathy: Implication in Diabetes ». Cardiovasc Res 109 (2): 217–27.

Chapouly, Candice, Sarah Guimbal, Pierre-Louis Hollier, et Marie-Ange Renault. 2019a. « Role of Hedgehog Signaling in Vasculature Development, Differentiation, and Maintenance ». International Journal of Molecular Sciences 20 (12). https://doi.org/10.3390/ijms20123076.

Chapouly, Candice, Sarah Guimbal, Pierre-Louis Hollier, et Marie-Ange Renault. 2019b. « Role of Hedgehog Signaling in Vasculature Development, Differentiation, and Maintenance ». International Journal of Molecular Sciences 20 (12). https://doi.org/10.3390/ijms20123076.

Chinchilla, P., L. Xiao, M. G. Kazanietz, et N. A. Riobo. 2010. « Hedgehog Proteins Activate Pro-Angiogenic Responses in Endothelial Cells through Non-Canonical Signaling Pathways ». Cell Cycle 9 (3): 570–79.

Hollier, Pierre-Louis, Candice Chapouly, Aissata Diop, Sarah Guimbal, Lauriane Cornuault, Alain-Pierre Gadeau, et Marie-Ange Renault. 2020. « Endothelial Cell Response to Hedgehog Ligands Depends on Their Processing ». BioRxiv, mars, 2020.03.03.974444. https://doi.org/10.1101/2020.03.03.974444.

Horng, S., A. Therattil, S. Moyon, A. Gordon, K. Kim, A. T. Argaw, Y. Hara, et al. 2017. « Astrocytic Tight Junctions Control Inflammatory CNS Lesion Pathogenesis ». J Clin Invest 127 (8): 3136–51. https://doi.org/10.1172/JCI91301 91301 [pii].

Jin, S., D. C. Martinelli, X. Zheng, M. Tessier-Lavigne, et C. M. Fan. 2015. « Gas1 Is a Receptor for Sonic Hedgehog to Repel Enteric Axons ». Proc Natl Acad Sci U S A 112 (1): E73–80. https://doi.org/10.1073/pnas.1418629112 1418629112 [pii].

Kenyon, B. M., E. E. Voest, C. C. Chen, E. Flynn, J. Folkman, et R. J. D’Amato. 1996. « A Model of Angiogenesis in the Mouse Cornea ». Invest Ophthalmol Vis Sci 37 (8): 1625–32.

Kogerman, Priit, Darren Krause, Fahimeh Rahnama, Lembi Kogerman, Anne Birgitte Undén, Peter G. Zaphiropoulos, et Rune Toftgård. 2002. « Alternative First Exons of PTCH1 Are Differentially Regulated in Vivo and May Confer Different Functions to the PTCH1 Protein ». Oncogene 21 (39): 6007–16. https://doi.org/10.1038/sj.onc.1205865.

Lee, C. S., L. Buttitta, et C. M. Fan. 2001a. « Evidence That the WNT-Inducible Growth Arrest-Specific Gene 1 Encodes an Antagonist of Sonic Hedgehog Signaling in the Somite ». Proc Natl Acad Sci U S A 98 (20): 11347–52.

Lee, C. S., L. Buttitta, et C. M. Fan. 2001b. « Evidence That the WNT-Inducible Growth Arrest-Specific Gene 1 Encodes an Antagonist of Sonic Hedgehog Signaling in the Somite ». Proceedings of the National Academy of Sciences of the United States of America 98 (20): 11347–52. https://doi.org/10.1073/pnas.201418298.

Li, Shuang, Shuangxi Li, Bing Wang, et Jin Jiang. 2018. « Hedgehog reciprocally controls trafficking of Smo and Ptc through the Smurf family of E3 ubiquitin ligases ». Science signaling 11 (516). https://doi.org/10.1126/scisignal.aan8660.

Martinelli, D. C., et C. M. Fan. 2007. « Gas1 Extends the Range of Hedgehog Action by Facilitating Its Signaling ». Genes Dev 21 (10): 1231–43.

Nie, D. M., Q. L. Wu, P. Zheng, P. Chen, R. Zhang, B. B. Li, J. Fang, L. H. Xia, et M. Hong. 2016. « Endothelial Microparticles Carrying Hedgehog-Interacting Protein Induce Continuous Endothelial Damage in the Pathogenesis of Acute Graft-versus-Host Disease ». Am J Physiol Cell Physiol 310 (10): C821–35. https://doi.org/10.1152/ajpcell.00372.2015 ajpcell.00372.2015 [pii].

Nusslein-Volhard, C., et E. Wieschaus. 1980. « Mutations Affecting Segment Number and Polarity in Drosophila ». Nature 287 (5785): 795–801.

Okada, A., F. Charron, S. Morin, D. S. Shin, K. Wong, P. J. Fabre, M. Tessier-Lavigne, et S. K. McConnell. 2006. « Boc Is a Receptor for Sonic Hedgehog in the Guidance of Commissural Axons ». Nature 444 (7117): 369–73.

Ramsbottom, Simon A., et Mary E. Pownall. 2016. « Regulation of Hedgehog Signalling Inside and Outside the Cell ». Journal of Developmental Biology 4 (3). https://doi.org/10.3390/jdb4030023.

Renault, M. A., J. Roncalli, J. Tongers, T. Thorne, E. Klyachko, S. Misener, O. V. Volpert, et al. 2010. « Sonic Hedgehog Induces Angiogenesis via Rho Kinase-Dependent Signaling in Endothelial Cells ». J Mol Cell Cardiol 49 (3): 490–98.

Robbins, D. J., D. L. Fei, et N. A. Riobo. 2012. « The Hedgehog Signal Transduction Network ». Sci Signal 5 (246): re6.

Sanchez-Arrones, Luisa, Marcos Cardozo, Francisco Nieto-Lopez, et Paola Bovolenta. 2012. « Cdon and Boc: Two Transmembrane Proteins Implicated in Cell-Cell Communication ». The International Journal of Biochemistry & Cell Biology 44 (5): 698–702. https://doi.org/10.1016/j.biocel.2012.01.019.

Sekiguchi, H., M. Ii, K. Jujo, M. A. Renault, T. Thorne, T. Clarke, A. Ito, et al. 2012. « Estradiol Triggers Sonic-hedgehog-induced Angiogenesis During Peripheral Nerve Regeneration by Downregulating Hedgehog-interacting Protein ». Lab Invest.

Tenzen, T., B. L. Allen, F. Cole, J. S. Kang, R. S. Krauss, et A. P. McMahon. 2006. « The Cell Surface Membrane Proteins Cdo and Boc Are Components and Targets of the Hedgehog Signaling Pathway and Feedback Network in Mice ». Dev Cell 10 (5): 647–56.

Xia, Y. P., Q. W. He, Y. N. Li, S. C. Chen, M. Huang, Y. Wang, Y. Gao, et al. 2013. « Recombinant Human Sonic Hedgehog Protein Regulates the Expression of ZO-1 and Occludin by Activating Angiopoietin-1 in Stroke Damage ». PLoS One 8 (7): e68891.

Zhang, Wei, Jong-Sun Kang, Francesca Cole, Min-Jeong Yi, et Robert S. Krauss. 2006. « Cdo Functions at Multiple Points in the Sonic Hedgehog Pathway, and Cdo-Deficient Mice Accurately Model Human Holoprosencephaly ». Developmental Cell 10 (5): 657–65. https://doi.org/10.1016/j.devcel.2006.04.005.

