## Supplementary data for "Desert Hedgehog-driven endothelium integrity is enhanced by Gas1 but negatively regulated by Cdon"

### Supplemental Material

**Supplemental figures and figure legends**

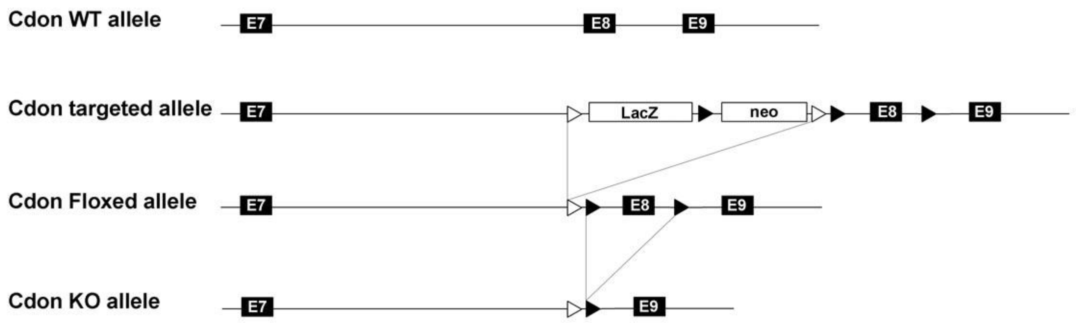


**Supplemental Figure 1:** Schema representing the Cdon Floxed allele.

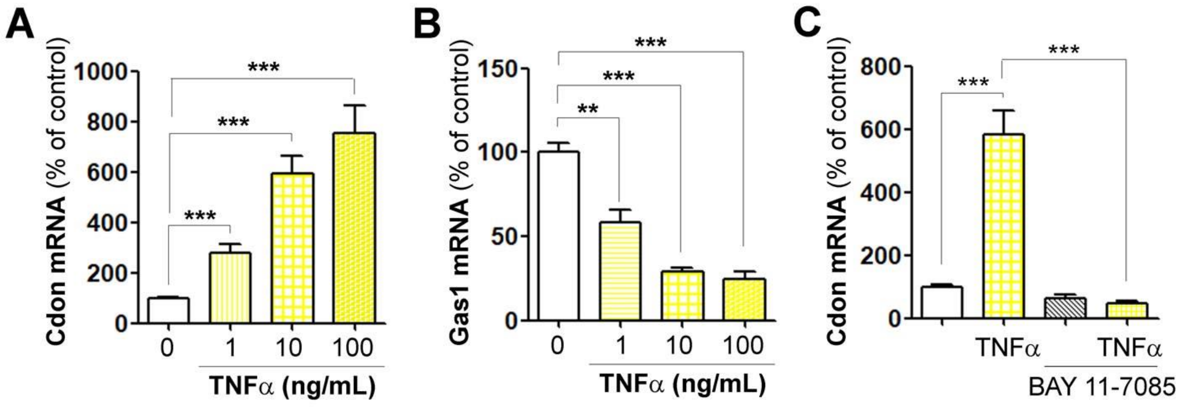


**Supplemental Figure 2: Cdon and Gas1 are regulated by TNFα in an opposite manner.** HUVECs were treated with 1, 10 and 100 ng/mL TNFα for 6 hours. Cdon (**A**) and Gas1 (**B**) mRNA expression was quantified via RT-qPCR. The experiment was repeated 3 times, each experiment included triplicates. (**C**) HUVECs were treated with 10 ng/mL TNFα in the presence or not of 2 μM BAY 11-7085 for 6 hours. Cdon mRNA expression was quantified via RT-qPCR. The experiment was repeated 3 times, each experiment included triplicates. **: $p \leq 0.01$; ***: $p \leq 0.001$. One way ANOVA followed by Bonferroni's multiple comparisons test.

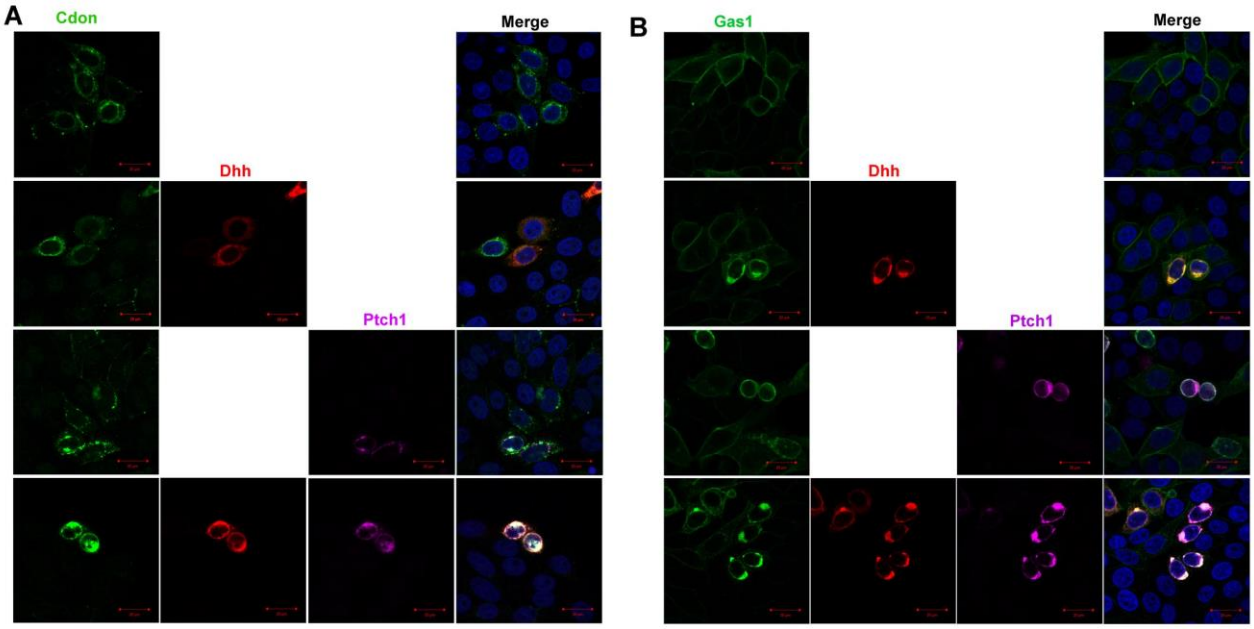


**Supplemental Figure 3:** (**A**) HeLa were co-transfected with Cdon, Dhh, and/or Ptch1 encoding plasmids. Cdon (in green), Dhh (in red) and Ptch1 (in purple) localization was assessed by immunostaining. (**B**) HeLa were co-transfected with Gas1, Dhh, and/or Ptch1 encoding plasmids. Gas1 (in green), Dhh (in red) and Ptch1 (in purple) localization was assessed by immunostaining.

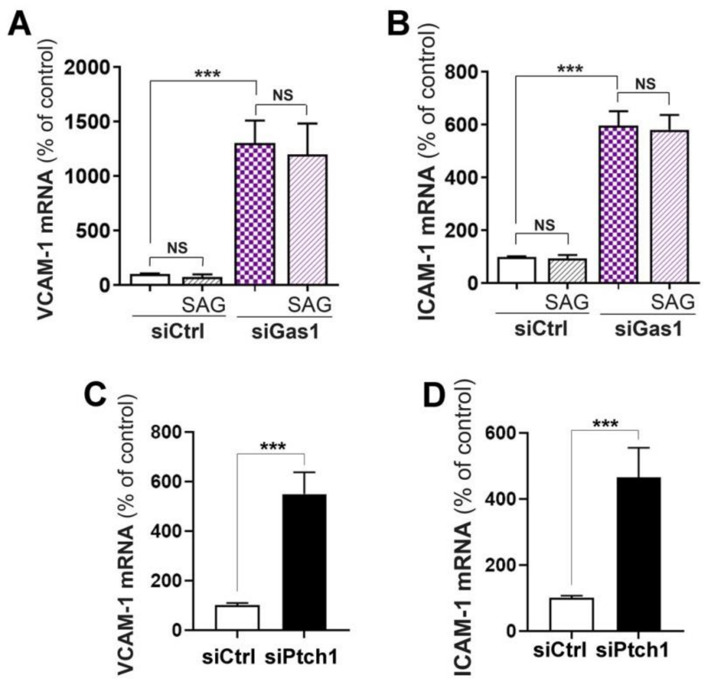


**Supplemental Figure 4: Ptch1 downregulates ICAM-1 and VCAM-1 mRNA independently on Smo.** (**A**-**B**) HUVECs were transfected with Gas1 or control siRNAs, then treated or not with 100 nM SAG. VCAM-1 (**A**) and ICAM-1 (**B**) mRNA expression was quantified via RT-qPCR. The experiment was repeated 3 times, each experiment included triplicates. (**C**-**D**) HUVECs were transfected with Ptch1 or control siRNAs. VCAM-1 (**C**) and ICAM-1 (**D**) mRNA expression was quantified via RT-qPCR. The experiment was repeated 3 times, each experiment included triplicates. **: $p \leq 0.01$; ***: $p \leq 0.001$. NS: not significant. Mann Whitney test or one way ANOVA followed by Bonferroni's multiple comparisons test.

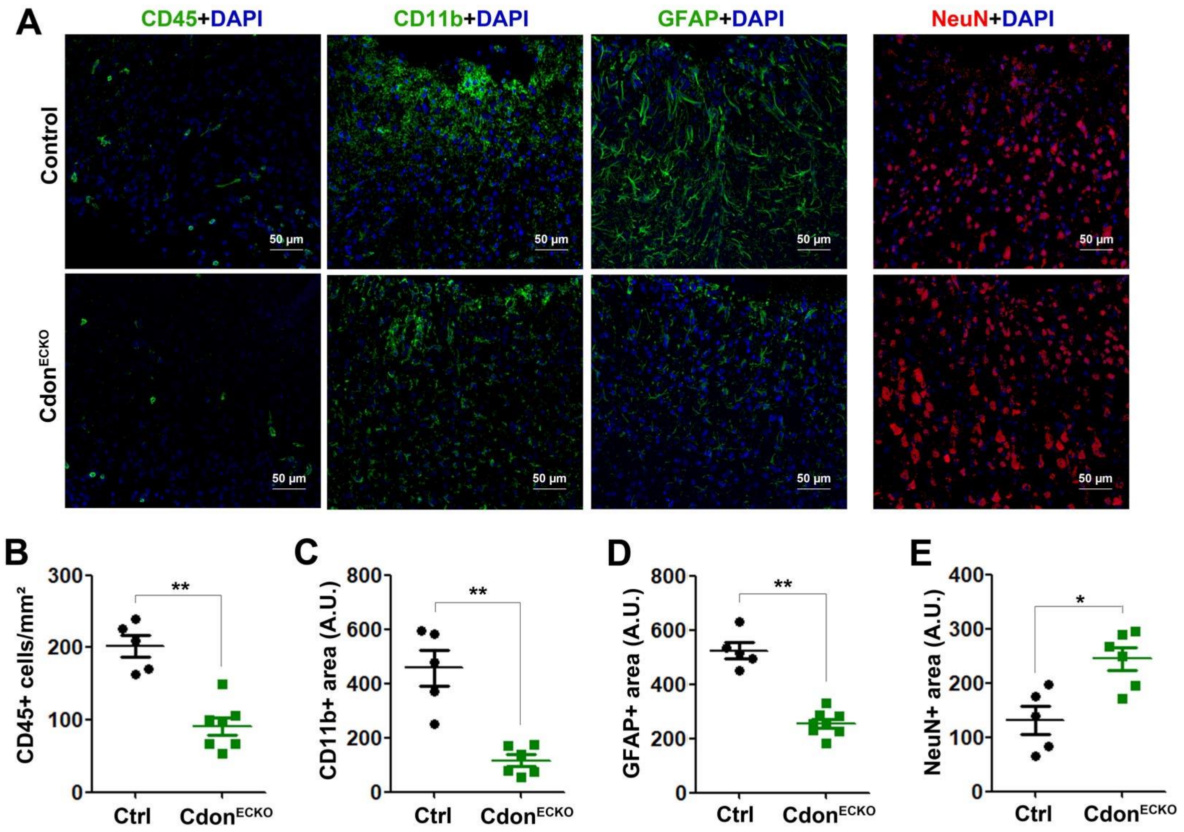


**Supplemental Figure 5:** Both Cdh5-Cre$^{ERT2}$; Cdon$^{Flox/Flox}$ (Cdon$^{ECKO}$) and Cdon$^{Flox/Flox}$ (control) mice were administered in the cerebral cortex with adenoviruses encoding Il1β (n=7 and 5 mice respectively). Mice were sacrificed 7 days later. (**A**) Brain sagittal sections were immunostained with anti-CD45 (in green), anti-CD11b (in green), anti-GFAP (in green) or anti-NeuN (in red) antibodies. Representative confocal images are shown. (**B**) Leucocyte infiltration was quantified as the number of CD45+ cells/mm². (**C**) Microglia activation was quantified as the CD11b+ surface area. (**D**) Reactive astrocytes were quantified as the GFAP+ surface area. (**E**) Neuronal survival was quantified as the NeuN+ surface area. *: p≤0.05; **: p≤0.01. Mann Withney test.

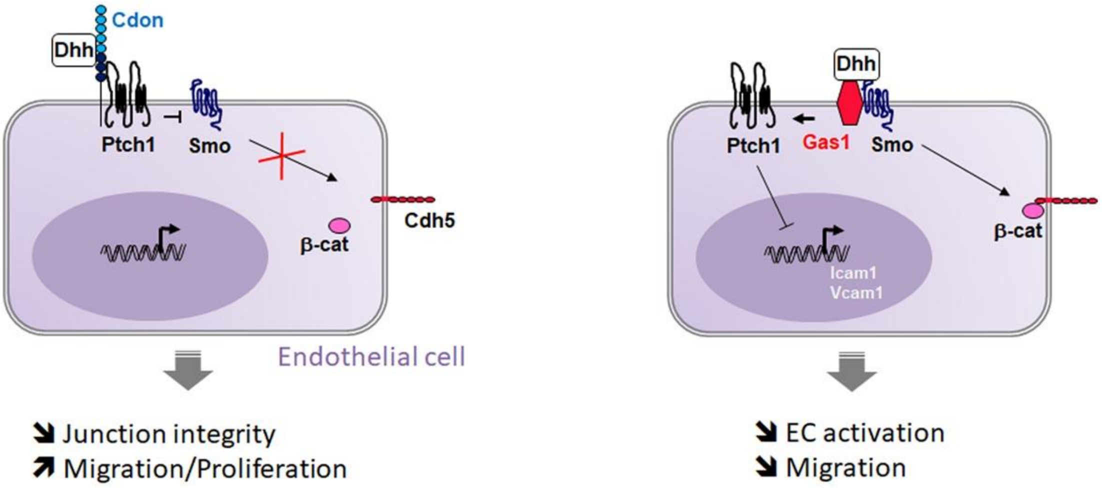


**Supplemental Figure 6: Schema recapitulating Gas1 and Cdon effects in ECs.**

**Supplemental Table**

| | | |
|---|---|---|
| hβ-actin | **F** | 5’-GGAGGAGCTGGAAGCAGCC-3’ |
| | R | 5’-GCTGTGCTACGTCGCCCTG-3’ |
| hCdon | F | 5’-TGGCCAGTTGCCGGAGGAGA-3’ |
| | R | 5’-ACAGCCCTCGGGGACAGGTG-3’ |
| hBoc | F | 5’-GACCCTCACCAGACTTGACC-3’ |
| | R | 5’-GGTCGTCTCTCTGGATCTGC-3’ |
| hGas1 | F | 5’-GCTAGCTGCAGTGTTCAGGA-3’ |
| | R | 5’-ATCCTCACTGGCCACAATCTG-3’ |
| hHhip | F | 5’-CTCGGCACTAGTGGGTCCTG-3’ |
| | R | 5’-TGCAGGTTGTACCGTGGCTC-3’ |
| hVCAM-1 | F | 5’-GGCCCAGTTGAAGGATGCGGG-3’ |
| | R | 5’-AGAGCACGAGAAGCTCAGGAGAA-3’ |
| hICAM-1 | F | 5’-ACGCCGGAGGACAGGGCATT-3’ |
| | R | 5’-GGGGCTATGTCTCCCCCACCA-3’ |

*F forward, R reverse*
*[illegible]-actin was used as the household gene*

**Supplemental Table I: List of primers used for reverse transcription (RT) quantitative polymer chain reaction (qPCR)**
